## Supplemental Table S1 for "Neural correlates of perceiving and interpreting engraved prehistoric patterns as human production: effect of archaeological expertise"

*Table S1 : Mean value and standard deviation of the BOLD signal in the 64 hROIs activated by at least one of the two groups of participants. in Attribution minus Orientation contrast.*

**BOLD signal in [Attribution *minus* Orientation]**

| | Controls | | | | Experts | | | |
|---|---|---|---|---|---|---|---|---|
| | Left hemisphere | | Right hemisphere | | Left hemisphere | | Right hemisphere | |
| | Mean BOLD | SD | Mean BOLD | SD | Mean BOLD | SD | Mean BOLD | SD |
| G_Cingulum_Ant-2 | - | - | 0.30 | 0.23 | - | - | 0.29 | 0.24 |
| G_Cingulum_Mid-2 | 0.15 | 0.23 | 0.21 | 0.25 | 0.23 | 0.16 | 0.24 | 0.17 |
| G_Cingulum_Post-1 | - | - | 0.15 | 0.21 | - | - | 0.05 | 0.30 |
| G_Cingulum_Post-3 | 0.08 | 0.29 | - | - | 0.39 | 0.36 | - | - |
| G_Frontal_Sup_Medial-3 | - | - | 0.20 | 0.20 | - | - | 0.26 | 0.25 |
| G_Fusiform-1 | 0.16 | 0.12 | - | - | 0.09 | 0.12 | - | - |
| G_Fusiform-2 | 0.20 | 0.17 | 0.14 | 0.16 | 0.11 | 0.13 | 0.06 | 0.09 |
| G_Fusiform-3 | 0.13 | 0.17 | - | - | 0.15 | 0.15 | - | - |
| G_Fusiform-4 | 0.29 | 0.36 | 0.16 | 0.22 | 0.28 | 0.21 | 0.19 | 0.15 |
| G_Fusiform-5 | 0.33 | 0.18 | 0.30 | 0.21 | 0.28 | 0.19 | 0.21 | 0.17 |
| G_Fusiform-6 | 0.30 | 0.24 | 0.32 | 0.24 | 0.25 | 0.19 | 0.21 | 0.17 |
| G_Fusiform-7 | 0.27 | 0.35 | - | - | 0.19 | 0.21 | - | - |
| G_Hippocampus-2 | 0.07 | 0.15 | 0.09 | 0.12 | 0.12 | 0.1 | 0.11 | 0.12 |
| G_Insula-anterior-2 | 0.36 | 0.32 | 0.31 | 0.23 | 0.34 | 0.2 | 0.33 | 0.22 |
| G_Insula-anterior-3 | 0.36 | 0.31 | 0.29 | 0.25 | 0.34 | 0.34 | 0.33 | 0.3 |
| G_Occipital_Lat-2 | 0.40 | 0.46 | 0.38 | 0.41 | 0.4 | 0.3 | 0.41 | 0.27 |
| G_Occipital_Lat-3 | 0.37 | 0.51 | 0.28 | 0.49 | 0.48 | 0.38 | 0.36 | 0.39 |
| G_Occipital_Lat-4 | 0.51 | 0.39 | 0.55 | 0.4 | 0.36 | 0.28 | 0.43 | 0.32 |
| G_Occipital_Lat-5 | 0.34 | 0.26 | 0.38 | 0.29 | 0.28 | 0.25 | 0.25 | 0.21 |
| G_Occipital_Mid-1 | 0.16 | 0.27 | 0.16 | 0.17 | 0.17 | 0.26 | 0.17 | 0.17 |
| G_Occipital_Mid-2 | 0.15 | 0.19 | 0.13 | 0.18 | 0.13 | 0.19 | 0.07 | 0.15 |
| G_Occipital_Pole-1 | 0.36 | 0.47 | 0.39 | 0.66 | 0.54 | 0.33 | 0.43 | 0.36 |
| G_ParaHippocampal-2 | - | - | 0.24 | 0.23 | - | - | 0.14 | 0.14 |
| G_Supp_Motor_Area-1 | 0.27 | 0.40 | 0.32 | 0.31 | 0.21 | 0.19 | 0.34 | 0.34 |
| N_Caudate-4 | 0.04 | 0.15 | - | - | 0.18 | 0.18 | - | - |
| N_Caudate-5 | 0.21 | 0.30 | 0.19 | 0.38 | 0.33 | 0.28 | 0.45 | 0.34 |
| N_Caudate-6 | - | - | 0.05 | 0.23 | - | - | 0.18 | 0.19 |
| N_Thalamus-1 | 0.26 | 0.36 | 0.21 | 0.41 | 0.58 | 0.36 | 0.57 | 0.43 |
| N_Thalamus-2 | -0.00 | 0.28 | 0.05 | 0.26 | 0.21 | 0.24 | 0.23 | 0.24 |
| N_Thalamus-3 | 0.29 | 0.44 | - | - | 0.38 | 0.32 | - | - |
| N_Thalamus-4 | 0.15 | 0.38 | 0.19 | 0.36 | 0.50 | 0.31 | 0.52 | 0.39 |
| N_Thalamus-5 | 0.08 | 0.19 | 0.12 | 0.2 | 0.09 | 0.13 | 0.09 | 0.14 |
| N_Thalamus-7 | - | - | 0.11 | 0.29 | - | - | 0.14 | 0.19 |
| S_Cingulate-1 | 0.34 | 0.32 | 0.39 | 0.34 | 0.28 | 0.24 | 0.34 | 0.27 |
| S_Cingulate-2 | 0.21 | 0.29 | 0.26 | 0.23 | 0.15 | 0.2 | 0.16 | 0.2 |
| S_Inf_Frontal-1 | - | - | 0.36 | 0.31 | - | - | 0.23 | 0.30 |
| S_Inf_Frontal-2 | - | - | 0.20 | 0.28 | - | - | 0.22 | 0.27 |
| S_Intraoccipital-1 | - | - | 0.31 | 0.32 | - | - | 0.27 | 0.32 |
| S_Orbital-1 | - | - | 0.15 | 0.21 | - | - | 0.10 | 0.15 |
| S_Orbital-2 | - | - | 0.46 | 0.33 | - | - | 0.47 | 0.2 |
| S_Precentral-4 | - | - | 0.06 | 0.18 | - | - | 0.09 | 0.14 |
