## Supplemental Table S2 for "Neural correlates of perceiving and interpreting engraved prehistoric patterns as human production: effect of archaeological expertise"

*Table S2: MNI coordinates of the 64 hROIs activated by at least one of the two groups of participants. in Attribution minus Orientation contrast.*

| hROIs in [Attribution - Orientation] | MNI coordinates (x y z) Left hemisphere | Right hemisphere |
|---|---|---|
| G_Cingulum_Ant-2 | - | 7 33 23 |
| G_Cingulum_Mid-2 | -4 3 30 | 4 4 30 |
| G_Cingulum_Post-1 | - | 5 -26 29 |
| G_Cingulum_Post-3 | -5 43 10 | - |
| G_Frontal_Sup_Medial-3 | - | 6 33 44 |
| G_Fusiform-1 | -2 -9 -34 | - |
| G_Fusiform-2 | -35 -26 -23 | 38 -25 -24 |
| G_Fusiform-3 | -37 -32 -24 | |
| G_Fusiform-4 | -43 -50 -17 | 44 -46 -18 |
| G_Fusiform-5 | -31 -50 -12 | 32 -47 -11 |
| G_Fusiform-6 | -28 -66 -10 | 29 -62 -9 |
| G_Fusiform-7 | -23 -84 -10 | - |
| G_Hippocampus-2 | -25 -32 -3 | 25 -31 -2 |
| G_Insula-anterior-2 | -34 17 -13 | 19 7 -19 |
| G_Insula-anterior-3 | -34 24 1 | 37 24 0 |
| G_Occipital_Lat-2 | -26 -94 -1 | 28 -89 -2 |
| G_Occipital_Lat-3 | -40 -84 -12 | 43 -81 -10 |
| G_Occipital_Lat-4 | -31 -89 8 | 34 -85 9 |
| G_Occipital_Lat-5 | -35 -79 -1 | 36 -76 2 |
| G_Occipital_Mid-1 | -32 -78 25 | 36 -74 25 |
| G_Occipital_Mid-2 | -37 -77 14 | 41 -73 12 |
| G_Occipital_Pole-1 | -21 -96 -14 | 24 -93 -13 |
| G_ParaHippocampal-2 | - | 29 -25 -19 |
| G_Supp_Motor_Area-1 | -6 22 46 | 6 21 48 |
| N_Caudate-4 | -15 11 12 | - |
| N_Caudate-5 | -13 10 8 | 12 10 9 |
| N_Caudate-6 | -16 4 19 | 15 7 18 |
| N_Thalamus-1 | -4 0 1 | 4 0 1 |
| N_Thalamus-2 | -9 -9 13 | 9 -7 13 |
| N_Thalamus-3 | -3 -7 -1 | - |
| N_Thalamus-4 | -3 -14 8 | 3 -14 9 |
| N_Thalamus-5 | -12 -19 7 | 13 -17 6 |
| N_Thalamus-7 | -9 -28 11 | 9 -26 10 |
| S_Cingulate-1 | -7 27 30 | 7 27 31 |
| S_Cingulate-2 | -7 16 41 | 8 14 46 |
| S_Inf_Frontal-1 | - | 46 40 10 |
| S_Inf_Frontal-2 | - | 44 19 28 |
| S_Intraoccipital-1 | - | 40 -40 51 |
| S_Orbital-1 | - | 25 41 -15 |
| S_Orbital-2 | - | 29 34 -13 |
| S_Precentral-4 | - | 44 1 48 |
